# Minimizing Structural Bias in Single-Molecule Super-Resolution Microscopy

**DOI:** 10.1101/293670

**Authors:** Hesam Mazidi, Jin Lu, Arye Nehorai, Matthew D. Lew

## Abstract

Single-molecule localization microscopy (SMLM) depends on sequential detection and localization of individual molecular blinking events. Due to the stochasticity of single-molecule blinking and the desire to improve SMLM’s temporal resolution, algorithms capable of analyzing frames with a high density (HD) of active molecules, or molecules whose images overlap, are a prerequisite for accurate location measurements. Thus far, HD algorithms are evaluated using scalar metrics, such as root-mean-square error, that fail to quantify the *structure* of errors caused by the *structure* of the sample. Here, we show that the spatial distribution of localization errors within super-resolved images of biological structures are vectorial in nature, leading to systematic structural biases that severely degrade image resolution. We further demonstrate that the shape of the microscope’s point-spread function (PSF) fundamentally affects the characteristics of imaging artifacts. We built a Robust Statistical Estimation algorithm (RoSE) to minimize these biases for arbitrary structures and PSFs. RoSE accomplishes this minimization by estimating the likelihood of blinking events to localize molecules more accurately and eliminate false localizations. Using RoSE, we measure the distance between crossing microtubules, quantify the morphology of and separation between vesicles, and obtain robust recovery using diverse 3D PSFs with unmatched accuracy compared to state-of-the-art algorithms.

## Introduction

Since its invention, fluorescence imaging has been an indispensable tool for biological studies of cells, tissues, and organisms because of its ability to visualize specific molecules of interest against a dark background in a relatively noninvasive manner. Tagging a biological molecule with a small organic fluorophore or fluorescent protein enables a fluorescence microscope to produce pictures of structures and movies of interactions between molecules within living cells. The optical detection of individual fluorescent molecules in condensed matter^1^ is the basis for an entire family of super-resolved fluorescence microscopy techniques^2–5^. These methods rely upon the blinking of fluorescent molecules in time to reduce the concentration of active emitters and resolve each molecule in a microscope image^6–8^. Repeated cycles of molecular blinking and measurement of molecular positions from their point spread functions (PSFs) by an image analysis algorithm result in reconstructed images of a biological structure with resolution beyond the Abbé diffraction limit (~ *;/2NA* ≎ 250 nm for visible light, where *NA* is the numerical aperture of the fluorescence microscope). Here, we refer to these techniques collectively as single-molecule localization microscopy (SMLM).

Although the experimenter often chooses imaging conditions to minimize the probability of image overlap between two molecules, the stochasticity of molecular blinking often leads to some overlap in SMLM datasets, especially for complex biological structures with high fluorophore labeling density^9^. One may even purposefully increase the density of active fluorescent probes in any given camera acquisition, such that images of neighboring molecules frequently or regularly overlap, in order to improve the temporal resolution of SMLM. Consequently, fewer imaging frames are needed to reconstruct a target structure, thereby leading to decreased phototoxicity as well as a reduction in motion-blur artifacts^10, 11^.

From a statistical perspective, super-resolution imaging in the presence of significant image overlap poses two major problems: (i) identifying the underlying molecules, and (ii) estimating their positions and brightnesses. Strategies for resolving overlapping molecules are primarily based on two aspects of prior knowledge: molecules are sparsely distributed in space, and they repeatedly and independently blink over time^12^. The first strategy recasts the estimation of molecular positions as a sparse recovery optimization problem, where a sparsity prior regulates the solution^13, 14^. The second approach exploits molecular emission characteristics (e.g., uncorrelated and repeated blinking events from various molecules) by applying higher-order statistics^15, 16^ or Bayesian analysis^17^ on the images of blinking molecules.

Extracting quantities from SMLM images, such as distances between structures or the size and shape of nanodomains, require molecular estimates with high *precision*, i.e., having a minimal spread around the mean, as well as high *accuracy*, i.e., having a mean with minimal deviation from the true value11,18. In low-density (LD) SMLM, analyzing repeated and isolated images of molecules guarantees statistically-unbiased estimates^19^. In particular, localization errors, i.e., the vectors obtained by taking the difference between true and estimated positions, are randomly distributed primarily due to Poisson shot noise. Consequently, the super-resolved images from a sufficiently large number of localizations are faithful representations of the ground truth, as long as systematic inaccuracies due to model PSF mismatch^11, 20, 21^, aberration^22^, and insufficient labeling^23^ are properly removed. These statistical results, however, no longer apply when analyzing high-density (HD) images. For example, standard LD algorithms, although capable of resolving two closely-spaced molecules, underestimate their separation distance^24^, suggesting that the localization errors are skewed toward neighboring molecules. This systematic bias gradually decreases as the separation distance becomes larger, achieving a minimum value beyond an algorithm-dependent distance^24^. In fact, the cumulative error between a super-resolution image reconstructed by a recovery algorithm and its ground truth scales exponentially with the regularity of molecular positions^25^ or how many molecules actively emit light within a diffraction-limited region at any point in time.

Because the structure of interest dictates the regularity in the position of labeling molecules, these theoretical results imply that the extent and characteristics of reconstruction errors fundamentally depend on the said structure. Put differently, accuracy is no longer a scalar quantity but a *vectorial* one. We refer to systematic inaccuracies caused by the structure of the sample as *structural bias* (see Supplementary Information, section 1 for a precise definition). Metrics used to evaluate SMLM algorithms, such as root-mean-square error (RMSE), detection rate^26^, and Pearson’s correlation^16^, collapse the reconstruction errors into a single number; moreover, visibility analysis only quantifies measurement precision, not accuracy^16^. These metrics, however, fail to fully characterize the vectorial nature of errors and ensuing biases. Errors caused by overlapping images become more severe in 3D SMLM, where 3D PSFs are larger than their 2D counterparts and are used to localize molecules over a larger domain^27^. More importantly, 3D super-resolution methods utilize diverse encoding mechanisms for depth, and a recovery algorithm that can be easily adapted to different PSFs is currently lacking.

Herein, we present a Robust Statistical Estimation algorithm, termed RoSE, for HD super-resolution imaging with arbitrary PSFs. In contrast to prior HD algorithms, RoSE jointly recovers molecular position and brightness through a novel optimization framework, allowing for estimating the likelihood that photons in the SMLM dataset arise from true molecular blinking events. By leveraging (i) the spatial sparsity in molecular positions to identify closely-spaced molecules; and (ii) the temporal statistics of molecular blinking events to eliminate false localizations, RoSE achieves robust recovery against arbitrary image overlap due to sample structure and PSF.

We apply RoSE to recover realistic, bio-inspired structures in 2D (microtubules and densely-packed vesicles^23^) *in silico*. Our analysis reveals a sample-dependent bias for HD imaging and shows that RoSE reduces these structural biases compared to state-of-the-art algorithms. Notably, our analysis shows that scalar metrics such as RMSE and visibility are insufficient in quantifying the sample-dependent errors present in super-resolution images. We also find that RoSE reduces these structural errors in the presence of experimental noise, by analyzing a SMLM reference dataset^26^. Finally, we show how the structure of both the sample and the 3D PSF itself fundamentally affect the systematic inaccuracies in reconstructing dense arrangements of nuclear pore complexes *in silico* and demonstrate the robustness of RoSE in adapting to various 3D PSFs, namely the double-helix^28^ and tetrapod^29, 30^ PSFs.

## Results

### Theory

#### A joint signal model

Localizing single molecules with overlapping images is in general a continuous recovery problem: molecular positions lie within a continuous range rather than a discrete set of points. Beyond localization, estimating the brightness of fluorescence bursts plays a critical role for measuring molecular orientation^31^.

To address these challenges, we propose a joint signal model in which we consider a single molecule as an isotropic point source and assume that within each frame, no two molecules emit within a certain neighborhood. (This model can be easily extended to include the emission anisotropy of dipole emitters by including molecular orientation^32^.) Thus, each molecule within an image can be mapped to a unique nearest discrete grid point, each associated with a brightness and a set of position gradients (Fig. 1(a)). This signal model allows us to explicitly decouple the number of photons that a molecule emits from its continuous position, thus providing a robust estimator of these quantities with sub-pixel accuracy (see Methods). In addition, these estimates can be exploited for temporal analysis, which is discarded by other signal models^13, 14^. We describe the main steps of RoSE below.

**Figure 1.**
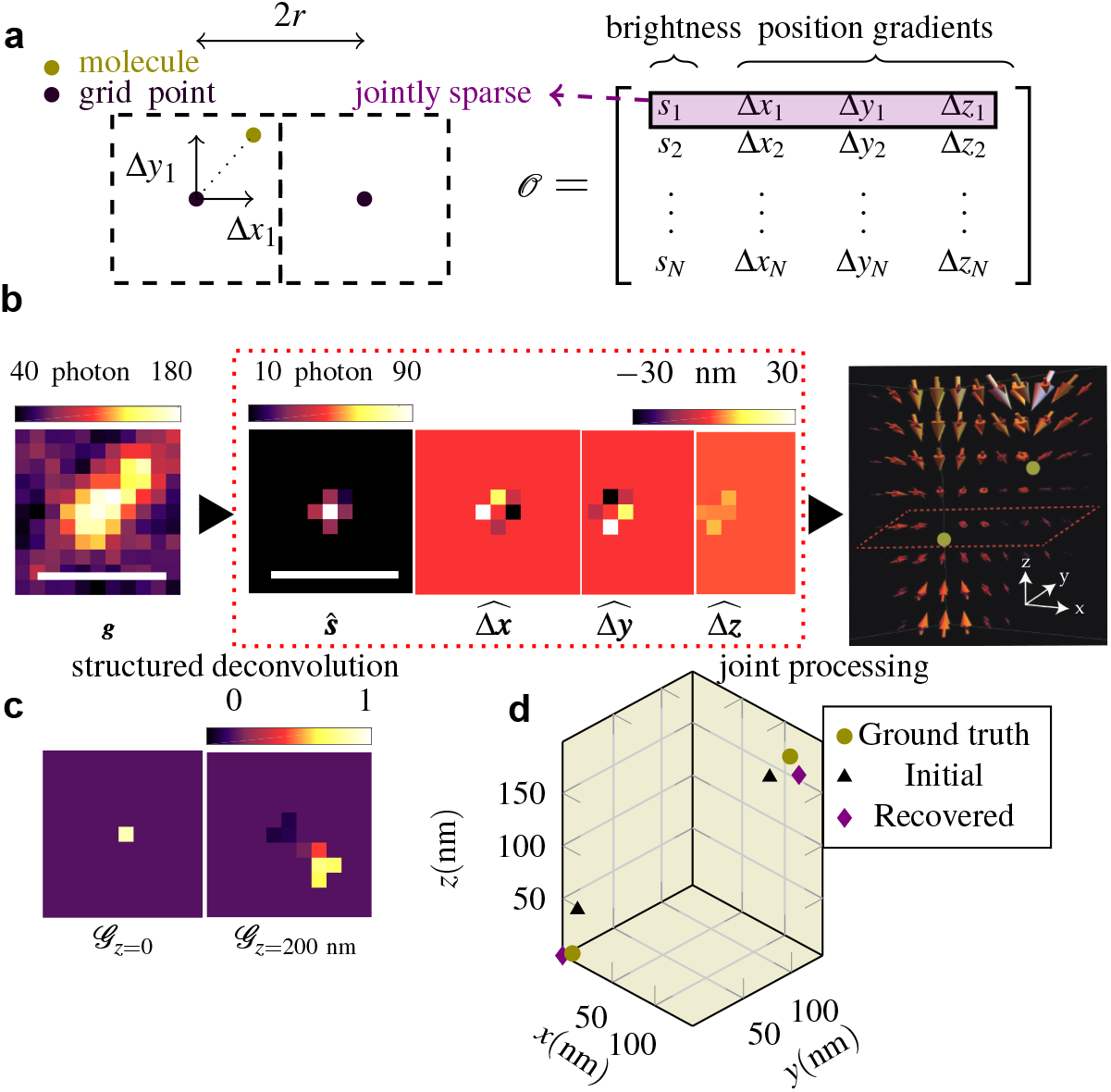
Joint recovery of molecular position and brightness by RoSE. (a) Left: Mapping of a continuous molecular position to a discrete grid in 2D (*r_x_ = r_y_ = r*). Right: Molecular parameters 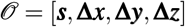 in 3D. (b) Left: Simulated image of two overlapping molecules, located at (0,0,0) and (120,120,180) nm using the tetrapod PSF. Middle: Slices of recovered parameters 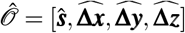 by (2) in the *z =* 0 plane. Right: Joint processing of brightness and position gradients of 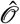 reveals two molecules unambiguously separated in 3D space (gold points). (c) Slices of estimated GradMap at *z =* 0 and z = 200 nm for the recovered signal in (b). Scale bars: 500 nm. (d) Initial position estimates (black triangles) obtained via the GradMap in (c), and the recovered molecular positions (purple diamonds) after applying adaptive constrained maximum likelihood. Ground-truth molecular positions are denoted by gold circles.

#### Identifying single molecules via structured deconvolution

When images of closely-spaced molecules overlap, the number of underlying molecules becomes an unknown parameter. The theory of sparse recovery provides algorithmic tools to identify single molecules from their overlapping images. Our approach leverages sparsity knowing that only a small number of molecules are active in each frame as compared to the number of grid points. This prior knowledge can be included in a deconvolution problem through a so-called regularizer such as *1_1_* norm while maintaining consistency with the measured data. Here, we propose a structured deconvolution method in which the regularizer enforces joint sparsity in molecular brightness and position, that is, if the brightness of a molecule associated with a grid point is zero (i.e., there is no molecule at that grid point), then the corresponding position gradients should be zero (see Methods).

Fig. 1(b) illustrates a 3D example of two closely-spaced molecules with significantly-overlapping images. The recovered molecular parameters from the structured deconvolution program denoted by 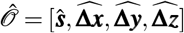 cannot resolve the brightnesses and positions of the two molecules. However, examining the joint structure of 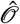 reveals that the brightness-weighted position gradients converge to the 3D positions of each molecule (Fig. 1(b), right). To make this mapping precise, we define a tensor *G*, called GradMap, in which each pixel takes on a value in [–1,1], termed the source coefficient, that signifies the local degree of convergence to that pixel (see Supplementary Information, section 3). In other words, the GradMap represents the likelihood that photons localized to particular positions in the map truly originate from the structure. Notably, GradMap leverages the convergent symmetry of the position gradients, thus sidestepping the need for PSF symmetry^16, 33^. Consequently, the number of molecules and their initial parameters are estimated from the local maxima of GradMap (Fig. 1(c)).

#### Accurate and precise recovery via adaptive maximum likelihood

After identifying the correct number of molecules, we further minimize the errors due to the sparsity constraint^14^ as well as the sample’s structure. Importantly, the errors from conventional sparse deconvolution programs, which may be larger than a few grid points, depends on the underlying molecular positions^25^. For accurate and precise estimates, RoSE adaptively updates the grid point closest to the current estimate of the molecule’s position (Fig. 1(d)) by maximizing adaptively a constrained maximum likelihood (see Supplementary Information, section 3). This strategy enhances the accuracy of both molecular position and brightness estimates and attains the limits of precision indicated by Cramér-Rao bound (CRB) (Figs. S2-S6).

#### Exploiting temporal blinking statistics via GradMap

Since molecular blinking events occur stochastically, misidentified localizations resulting from overlapping images exhibit significantly less autocorrelation over time compared to molecules that are accurately identified. Thus, the autocorrelated GradMap represents the confidence that molecules represent the true structure and not a false localization (see Supplementary Information, section 3). Noting that GradMap localizes these blinking events with sub-pixel accuracy, we apply a threshold on the pixel-wise temporal autocorrelation of GradMap to eliminate false localizations. We refer to this scheme as RoSE-C.

### Beyond Scalar Metrics

Traditional performance metrics for evaluating HD recovery algorithms, such as recall rate (i.e., the number of molecules that are recovered correctly with a certain distance criteria) and the RMSE between correctly recovered molecules and their matched ground truths^26^, disregard the role of underlying structure; the magnitude and direction of errors vary across the super-resolved image. In the following sections, we examine and characterize the structural biases of various recovery algorithms. Our analysis includes simulated realistic, HD SMLM image stacks of several bio-inspired structures as well as experimental SMLM images of microtubules.

#### Accurate frame-by-frame analysis: imaging crossing microtubules using RoSE

To quantitatively characterize the effects of sample structure and blinking density on the size and structure of localization errors, we construct and examine two crossing microtubules (MTs) *in silico*. MTs are tubular intracellular assemblies of the protein tubulin that form a dynamic, three-dimensional network^34^. MTs act as “robotic arms” and are responsible for key functions such as cell division, shaping the cell membrane, and intracellular transport^34^.

The MTs are simulated as cylinders aligned perpendicular to the optical axis with radii of 12.5 nm and are displaced symmetrically about the focal plane with an axial separation of 25 nm. We develop a stochastic model to generate SMLM images (see Supplementary Information, section 4). To measure the mean separation between MTs, we consider a sliding rectangular area containing both MTs with a constant 58.5-nm height (Fig. 2(a) orange box). The rectangle is sufficiently large such that labeling noise is insignificant but small enough to resolve algorithm performance as a function of true separation *d*\*. The localizations within the rectangular area are then projected onto the *x*-axis transverse to the MTs, collapsing the localization data into histograms.

**Figure 2.**
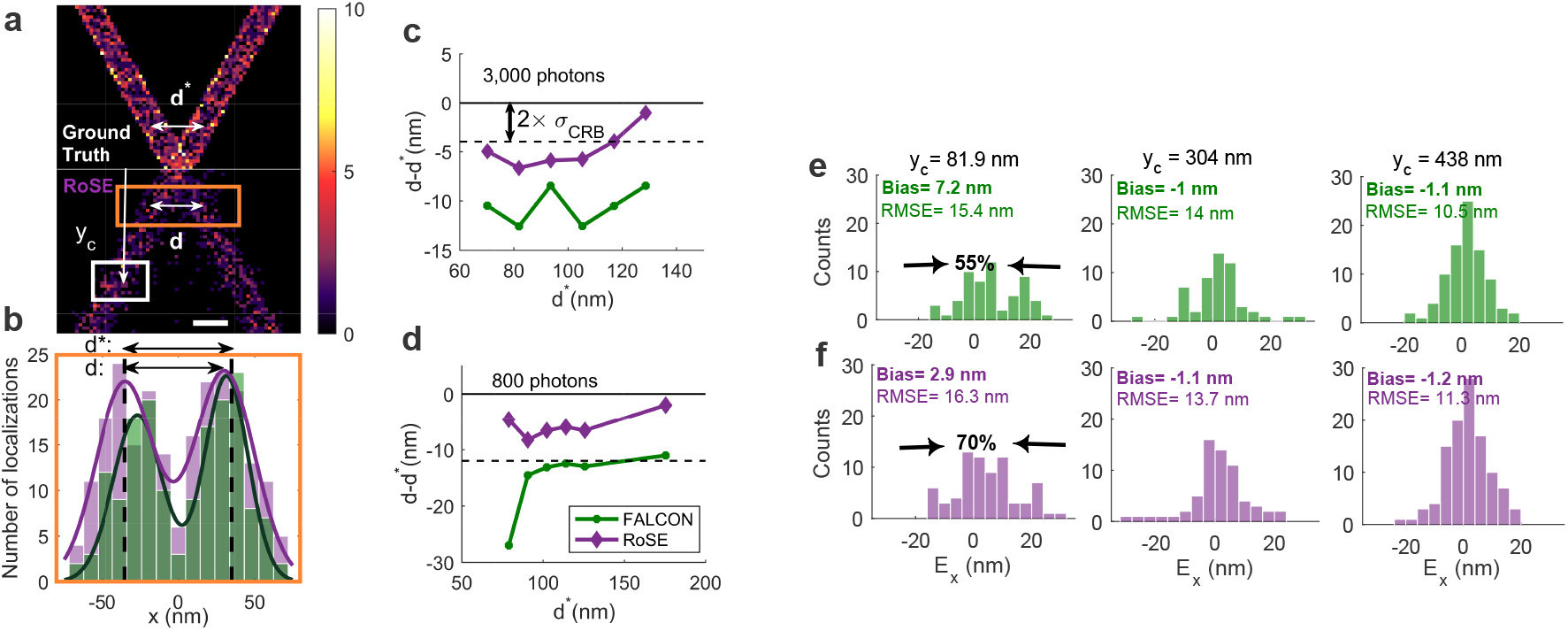
Structural bias of two crossing microtubules (MTs) recovered by RoSE (purple) and FALCON (green). (a) 2D histogram of (top) the simulated ground-truth structure and (bottom) recovered structural obtained by RoSE for a blinking density of 1.3 × 10^−5^ molecules/nm^2^. Scale bar: 50 nm. Color bar: number of localizations per 5 × 5 nm^2^. (b) Projection of the localizations within the orange box in (a) and the corresponding double Gaussian fits. The arrows labeled by d* and *d* denote the distance between the centers of the two MTs, and the distance between the peaks of the fitted double Gaussian, respectively. (c,d) Mean distance between centers of the MTs along the length of the structure for mean emission intensities of (c) 3000 photons and (d) 800 photons. (e,f) Localization errors along the +x direction *(E*_x_) for molecules within the white box in (a) at various distances from the center crossing y_c_: (left) 81.9 nm, (middle) 304 nm, (right) 438 nm. The noted bias values represent the difference between the true center of the MT and the mean of matched localizations; the arrows indicate a [10, −10] nm interval centered at the true position of the MT.

We tested RoSE and FALCON^14^, of which the latter has a high accuracy score (low RMSE) and a competitive Jaccard index^26^, on image data generated at various blinking densities, namely 4.9 × 10^−6^, 1.3 × 10^−5^, and 1.9 × 10^−5^ molecules/nm^2^, representing low, high, and ultra-high densities, respectively (Fig. S7). At high blinking density, the measured mean separation distance exhibits a negative bias for both algorithms, meaning errors are structured toward the center of the MTs (Fig. 2(b)). Compared to FALCON, RoSE reduces the bias in measuring *d\** across the entire length of the structure, and the measured errors drop to being CRB-limited more quickly for RoSE than FALCON for both low (800 photons) and high (3000 photons) molecular brightnesses (Fig. 2(c),(d)). In particular, for true mean separations from 70.2 nm to 128.7 nm (Fig. 2(c)), RoSE incurs an average bias of –4.7 nm compared to −10.5 nm for FALCON. This difference in average bias becomes more pronounced for dimmer molecules (Fig. 2(d)), where RoSE exhibits a small bias of −5.7 nm, which is well within the FWHM of the theoretical limit of precision 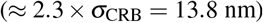. In contrast, FALCON incurs an average bias of −15.2 nm. For large *d*\*, the error distance is non-zero due to non-uniform sampling and stochastic localization precision.

Examining the left-most MT at various distances *y_c_* from the crossing point reveals the vectorial nature of these errors (Fig. 2(e,f)). We calculated the localization error *E_x_* along the x dimension by matching correctly-identified localizations to the ground-truth dataset^26^ (see Supplementary Information, section 5). Interestingly when *y_c_ =* 81.9 nm, the errors are skewed toward the +x direction (Fig. 2(e)), in accord with the systematic underestimates of MT separation (Fig. 2(c,d)). As *y_c_* increases, the errors converge to a Gaussian distribution centered at zero whose width is determined by the CRB, as expected.

RMSE analysis cannot reveal such a structural bias, i.e., the systematic deviation of localized molecules away from the true center of the MT. Surprisingly, the RMSE for FALCON is slightly smaller than RoSE for y_c_ = 81.9 nm (Fig. 2(e,f) left); however, the percentage of localized molecules for RoSE that are within [−10,10] nm is 70% compared to 55% for FALCON; this increased localization accuracy results in more accurate measurements of the MT separation distance (Fig. 2(b)). Indeed, the Jaccard index calculated over the entire structure for RoSE (54%) is higher than FALCON (49%). These results suggest that decomposing the reconstruction errors into scalar detection and localization error metrics cannot quantify the variation in magnitudes as well as vectorial nature of these significant biases. Therefore, a structure-based analysis of errors beyond scalar metrics is critical for quantitative evaluation and insight into the performance of recovery algorithms.

Additionally, RoSE maintains its robust performance and consistently outperforms FALCON at both low and higher blinking densities. Even at low blinking density, RoSE resolves the walls of MTs with much better visibility than FALCON (57% larger), especially close to the crossing point (Fig. S7).

#### Leveraging blinking statistics across frames: accurate imaging of vesicles via RoSE-C

SMLM techniques have revealed the morphology of protein clusters within bacteria^35^ and during the assembly of HIV viruses^36^. In particular, clustering algorithms have been utilized to infer clusters and their spatial distributions from SMLM datasets^18^. The accuracy of such analyses, however, critically relies upon the accuracy of positions output from the SMLM algorithm.

We examine the structural errors in characterizing clusters from simulated SMLM data of a dense arrangement of fluorescently-labeled vesicles. These four vesicles are modeled as circles with a 30 nm radius, each uniformly labeled with 100 molecules. These molecules blink within a total of 80 frames (a mean blinking density of 4.7 × 10^−5^ molecules/nm^2^), resulting in highly-overlapped images and a short acquisition time. To separate Poisson shot noise and sampling artifacts, we compared reconstructions from this HD dataset to equivalent LD imaging (i.e., one burst per frame with the same brightness as in HD data) in which no image overlaps. ThunderSTORM was chosen to provide a super-resolved structure from LD frames^37^.

We tested the performance of RoSE-C in eliminating false localizations that mostly occur between vesicles due to significant image overlap. Remarkably, RoSE-C perfectly restores the discrete structure of packed vesicles for various signal-to-background levels as well as blinking densities (Fig. S8). Computing the confidence level of obtaining true localizations, quantified by the auto-correlated GradMap, critically enables RoSE-C to filter out false localizations otherwise recovered by RoSE.

ThunderSTORM, FALCON, and RoSE-C are all able to resolve 4 clusters (Fig. 3(a-d)). The sizes and shapes of these vesicles were analyzed using a density-based clustering algorithm, which takes into account vesicle size and localization precision and is robust against false localizations^38^ (see Supplementary Information, section 5). The two left-most clusters for ThunderSTORM (Fig. 3(b,e)) exhibit a small average ellipticity of 1.2, defined by the ratio of the major axis and the minor axis, and similarly the two right-most clusters (Fig. 3(b,f)) have an average ellipticity of 1.12. These small ellipticity values could be caused by Poisson shot noise and finite-sampling artifacts. Interestingly, the clusters for both RoSE-C and FALCON (Fig. 3(c-f)) are skewed toward neighboring vesicles, and exhibit larger ellipticity values. The two left-most clusters recovered by RoSE-C (Fig. 3(d,e)) exhibit an average ellipticity of 1.38, while for FALCON (Fig. 3(c,e)), the average ellipticity is 1.84. For the two right-most clusters, RoSE-C (Fig. 3(d,f)) has a slightly smaller average ellipticity of 1.42 compared to FALCON’s (Fig. 3(c,f)) 1.44. Overall RoSE-C minimizes the average ellipticity bias by 24%. This analysis reveals vectorial inaccuracies (i.e., structured ellipticity) present in the super-resolved images that cannot be fully quantified by scalar RMSE values.

**Figure 3.**
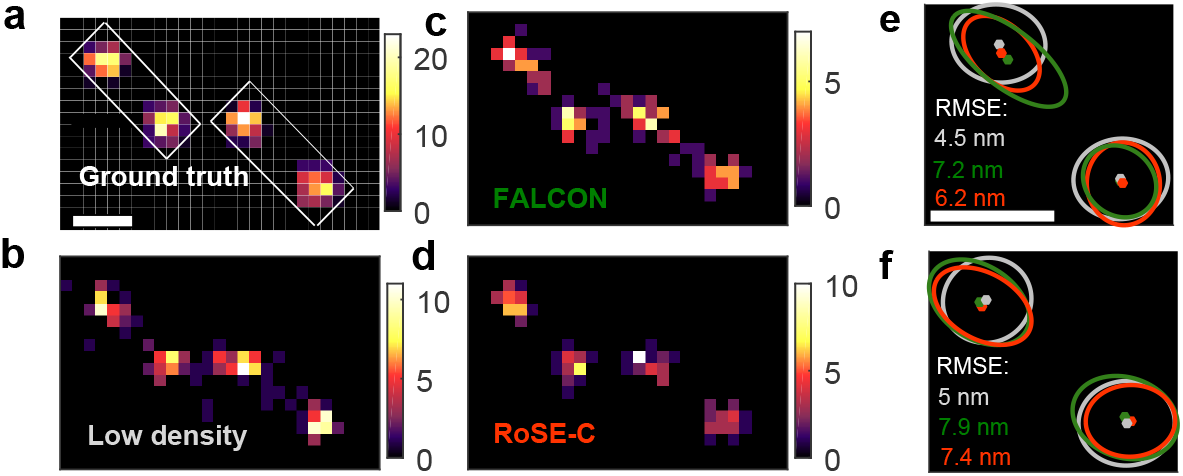
Structural bias in the sizes and shapes of vesicles recovered by RoSE-C and FALCON. (a) Simulated ground-truth structure of 4 circular vesicles. The mean emission intensity and mean uniform background were set to 800 photons and 40 photons per pixel, respectively. (b) Structure recovered from low-density frames using ThunderSTORM. (c) Structure recovered by FALCON. (d) Structure recovered by RoSE-C. Color bars: number of localizations per 19.5 × 19.5 nm^2^. (e,f) Estimated clusters, their confidence ellipses, and the RMSE of all localizations corresponding to the (e) left and (f) right boxes in (a) for low-density imaging (grey), FALCON (green), and RoSE-C (orange), respectively. Scale bars: 100 nm.

Recently, a class of algorithms termed super-resolution radial fluctuations (SRRF)^16^ was introduced based on superresolution optical fluctuation imaging (SOFI)^15^. SRRF relies on radial symmetry (spatial information) to reduce the apparent FWHM of the Gaussian PSF. Moreover, by applying higher-order statistics to images of blinking molecules over time, SRRF resolves molecules at smaller separations with better visibility. Analysis of simulated SMLM datasets using SRRF and RoSE-C reveals that SRRF images contain a vesicle separation bias of 27%, while RoSE-C reduces this bias to −3% (see Supplementary Information section 7, Figs. S9-S11). Visibility metrics fail to capture this bias.

### Structured errors in experimental SMLM images

Quantifying the structured bias present in SMLM images can be difficult without knowledge of the ground truth. However, microtubules have often been used in demonstrations of SMLM because of their smoothness and well-characterized branching behavior. We compare the performance of FALCON, SRRF, and RoSE on experimental SMLM images of MTs^26^ in regions where MT bundles are dense and the fluorescence background is relatively uniform. We observe that the cumulative errors in molecule position exponentially increase with molecular density in crowded regions (white rectangular region in Fig. 4(g-i)).

**Figure 4.**
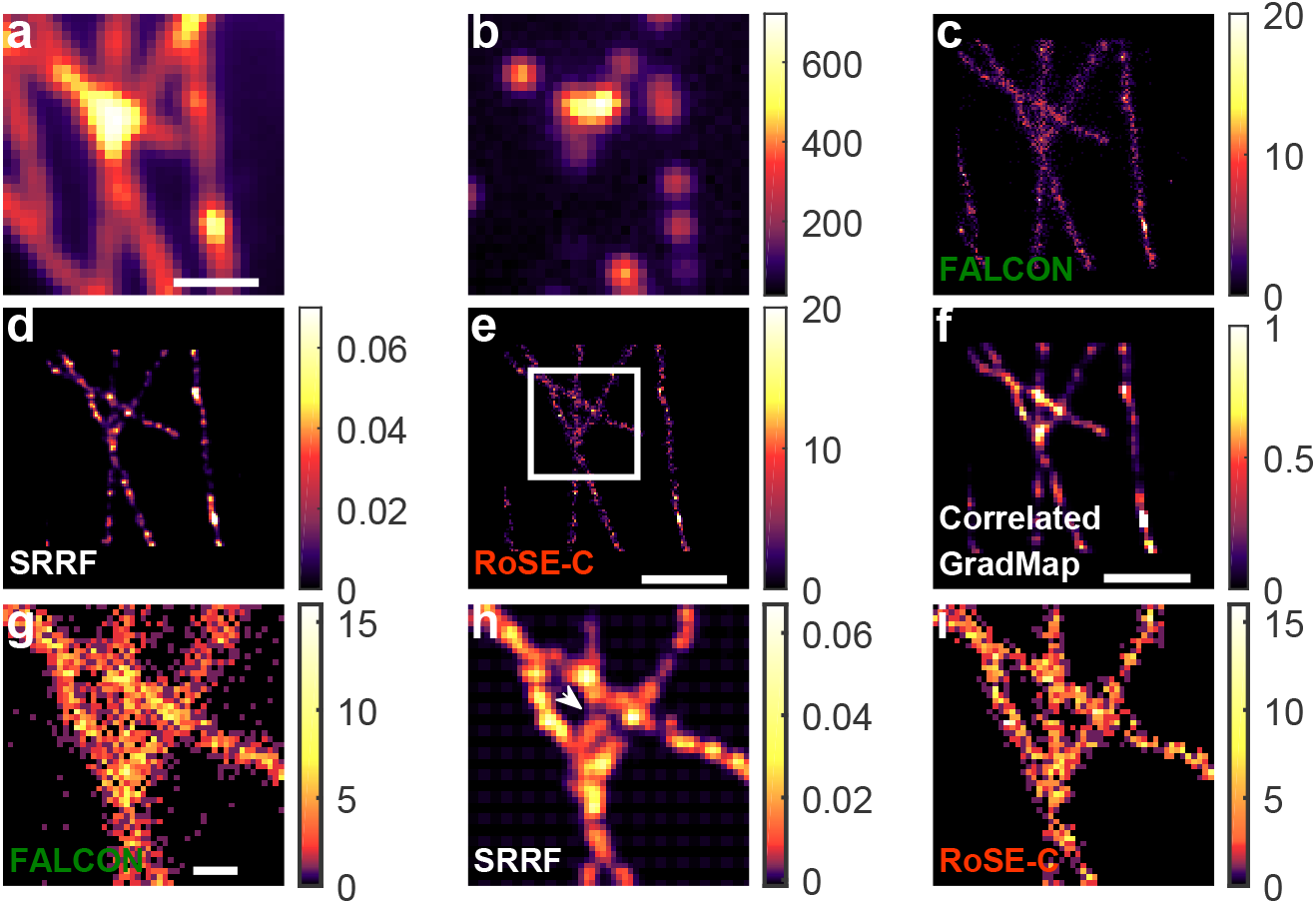
Structural bias in recovering a dense network of microtubules from experimental SMLM images using FALCON, SRRF and RoSE-C. (a) Diffraction-limited sum of the SMLM image stack. (b) An example SMLM image exhibiting high blinking density. (c) Histogram of localizations obtained using FALCON. (d) Recovered map using SRRF-TRAC2. (e) Histogram of localizations obtained using RoSE-C. (f) Correlated GradMap recovered using RoSE-C. The white arrow indicates regions with low confidence. (g-i) Magnified views of the boxed region in (e) for (g) FALCON, (h) SRRF, and (i) RoSE-C. The white arrow in (h) indicates a bias in the recovered branching structure. Scale bars: (a,e,f) 1 μm and (g) 200 nm. Color bars: (b) number of photons per 100 × 100 nm^2^ and (c,e,g,i) number of localizations per 25 × 25 nm^2^.

The nature of these errors depends on the recovery algorithm. The localizations obtained using FALCON generally broaden and occlude details of the branching structure (Fig. 4(g)). In contrast, the structure recovered by SRRF exhibits distortion of the middle MT such that it appears to extend towards the upper-right branch (Fig. 4(h)). RoSE-C appears to recover the structure of the MT network with both greater accuracy and precision (Fig. 4(i)). We note that the correlated GradMap exhibits low confidence for regions that do not belong to the MT network (Fig. 4(f)), which is consistent with the results of recovering densely-packed vesicles (Fig. 3).

We further consider another region in the dataset exhibiting non-uniform molecular blinking (Fig. S13(a)). For each algorithm, we observe structured bias in the SMLM reconstructions that are similar to the aforementioned simulation studies (Fig. 2): the separation distances between MTs measured using localizations recovered by FALCON are smaller than those obtained by RoSE-C (Fig. S13(e,f)). In addition, separation distances across the length of MTs for RoSE-C exhibit smooth curvature that is consistent with our expectation of MT network structure. On the other hand, separation distances obtained by FALCON exhibit significant biases where parallel MTs are closer together (Fig. S14(a,b,c)). As the MTs become far apart, the separation distances measured by each algorithm converge to similar values (Fig. S14(d)). Examining the recovered images by SRRF algorithms reveals the structured biases, wherein parallel MTs are frequently unresolved and visibility is difficult to quantify (Fig. S15).

The GradMap provides an insightful map of confidence levels, in which it displays low confidence for regions that are subject to localization errors due to either low sampling or dense molecular blinking (Fig. S13(d)). This confidence metric cannot be inferred from the SMLM reconstructions themselves produced by FALCON, SRRF, and RoSE-C. We note that in analyzing experimental SMLM images using RoSE-C, we chose the confidence threshold adaptively (see Supplementary Information, section 3. D), but all other parameters remained the same from the aforementioned *in silico* studies. We could not recover structures with similar quality as RoSE-C by tuning the sparsity parameter of FALCON or related parameters of SRRF (Fig. S16).

### Robust 3D recovery using diverse PSFs

PSF engineering has become a popular strategy for 3D super-resolution imaging. A variety of PSFs, including the astigmatic PSF^39^, the double-helix PSF (DH-PSF)^28^, and the tetrapod PSF^29, 30^ can be implemented with fixed phase masks or programmable hardware. These PSFs have tunable axial ranges (spanning 2 – 20 μm^27^) and exhibit diverse and complex features over their measurement domains. Importantly, they occupy more space on the camera compared to the standard PSF. Therefore, addressing the problem of image overlap for an arbitrary 3D PSF becomes critical.

We test the robustness of RoSE-C in recovering dense arrangements of nuclear pore complexes (NPCs) in 3D using the DH and tetrapod PSFs. An NPC is a ring-like protein complex composed of eight subunits with an approximate diameter of 90-120 nm, well below the diffraction limit^40^. The complex and dense 3D arrangements of NPCs necessitate the use of 3D super-resolution techniques for measuring their diameters and resolving their subunits^41^. We simulated each NPC using 8 subunits equidistantly distributed on a circle with a diameter of 120 nm. The NPCs are placed in 3D with centers at (0,0,0), (300,0,50), and (200,200,130) nm with various orientations (Fig. 5(a,b)). Each subunit is labeled with a single fluorophore that blinks stochastically with a mean brightness of 3100 photons per blinking event. On average, one molecule per NPC emits during any specific frame, causing a high probability of PSF overlap on the camera (Fig. 5(c,d)).

**Figure 5.**
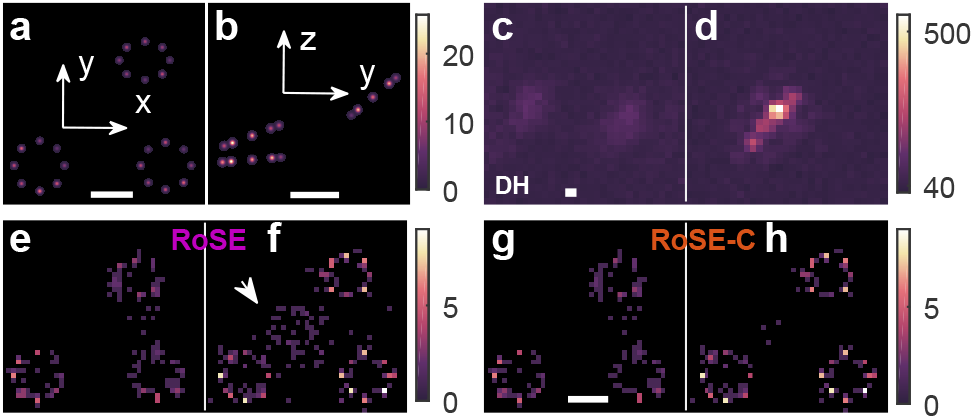
3D recovery of densely-packed NPCs using the double-helix and tetrapod PSFs. (a) xy and (b) yz views of the simulated ground-truth arrangement of 3 NPCs centered at (0,0,0), (300,0,50), and (200,200,130) nm, each consisting of 8 labeling sites equidistantly distributed on a circle with diameter of 120 nm. The brightest pixel corresponds to the peak of a Gaussian distribution (standard deviation = 3.5 nm) multiplied by the number of blinking events. (c,d) Representative images of overlapping molecules corresponding to the (c) DH and (d) tetrapod PSFs. (e,f) xy projections of the NPCs recovered by RoSE using the (e) DH and (f) tetrapod PSFs. The white arrow in (f) indicates mislocalizations at higher axial coordinates. (g,h) xy projections of the NPCs recovered by RoSE-C using the (g) DH and (h) tetrapod PSFs. Scale bars: 100 nm. Color bars: (a,b) number of blinking events/nm^2^; (c,d) number of photons per 58.5 × 58.5 nm^2^; and (e-h) number of localizations per 12 × 12 nm^2^.

In the projected 2D images of the NPCs recovered by RoSE, the NPC subunits are more blurry when using the DH-PSF (Fig. 5(e)) compared to the tetrapod (Fig. 5(f)). This difference is consistent with the superior precision of the tetrapod PSF^29^, which allows for counting subunits using the clustering algorithm with 25% better accuracy (see Fig. S17 for the performance of the DH and tetrapod PSFs in resolving two closely-spaced molecules). Intriguingly, however, the structure resolved by RoSE using the tetrapod PSF exhibits a set of mislocalizations at higher axial coordinates that resemble an NPC (Fig. 5(f)). In contrast, there are fewer mislocalizations in the reconstruction using the DH-PSF, and interestingly, they do not resemble an NPC (Fig. 5(e)).

RoSE-C remarkably minimizes the structured mislocalizations when using the tetrapod PSF (Fig. 5(h)). This improved accuracy is enabled by GradMap, in which pixels that correspond to false localizations exhibit smaller autocorrelation over time than those that correspond to true localizations (Fig. S18). For the DH-PSF, the mislocalizations between the two right-most NPCs cannot be eliminated with RoSE-C. One possible reason is that the pixel size of the GradMap (58.5 nm) is larger than the half of the separation between the two right-most NPCs along the y dimension (120 nm). Thus, in this particular simulation, GradMap has insufficient spatial sampling to resolve false localizations from true ones using temporal autocorrelation.

To gather insight into the structured mislocalizations in the recovered NPCs using the tetrapod PSF, we examined a different arrangement with NPCs located at (0,0,0), (300,0,50), and (200,200, −130) nm using the tetrapod PSF. Surprisingly, the number of mislocalizations in the super-resolved structure by RoSE is significantly reduced (Fig. S18). This reduction likely results from the distinct depth encoding of negative z values compared to positive ones for the tetrapod PSF. Thus, the tetrapod can resolve NPCs accurately when they are alternate sides of the focal plane rather than all above the focal plane. These results reveal the role of both structure and the degeneracy of PSF in causing biases (see Supplementary Information, section 8).

## Discussion

The *vectorial* nature of localization errors in SMLM is often ignored. Our simulations show that when recovering bio-inspired structures under HD imaging conditions, reconstruction errors exhibit magnitudes and directions that depend on the structure of the sample and cannot be revealed by conventional error metrics. Our analysis of HD experimental data of dense microtubules corroborates these results. Furthermore, in 3D SMLM, the inaccuracies are affected both by the *structure* of the sample and the *structure* of the PSF. In particular, PSF degeneracy causes structured mislocalizations that may pose difficulty for downstream quantitative analysis^18^.

Current methods for minimizing nanoscale inaccuracies, including classifying localizations based on spot size, brightness^24^, and similarity-based techniques^42^, lack robustness against sample structure and become unreliable for 3D imaging. Further, point-wise precision for individual localizations, which is used for filtering localizations with large uncertainty, becomes inaccurate in the case of overlapping PSFs due to biased brightness estimates (Fig. S4). Taken together, these observations necessitate estimation of a fundamentally different quantity for individual localizations.

RoSE addresses these challenges by directly estimating the likelihood of fluorescence emission for individual blinking events, which enables molecules with overlapping images to be localized in the continuous domain and increases localization and photon-counting accuracy. Further by assuming that the sample is static over the imaging interval, RoSE-C simply autocorrelates these likelihoods to obtain point-wise confidence levels without knowing the sample structure *a priori*. The confidence level of a localization signifies its uncertainty in representing the ground truth, which is exploited to eliminate false localizations at higher blinking densities (Fig. S8). More importantly, the estimated confidence levels do not rely on point-wise precision, brightness, or how molecules’ images overlap and can be applied to any arbitrary PSF. Access to such a confidence metric is a key to automating challenging quantitative analyses in SMLM, such as elucidating protein organization in 3D.

We emphasize that although RoSE-C eliminates localizations with small confidence levels, which may degrade the overall apparent labeling density, these localizations would otherwise lead to structural biases if they are not eliminated. This degradation in HD SMLM is fundamentally different from low-density SMLM for which even localizations with large uncertainty provide unbiased information regarding the underlying structure. Besides thresholding, the confidence levels can systematically be used as additional input to increase the accuracy of various quantitative analyses such as Bayesian clustering^43^. Nonetheless, RoSE-C exhibits robustness in measuring the apparent labeling density w.r.t. the number of blinking events per molecule, whereas image-based HD algorithms utilizing higher-order statistics incur significant errors (Fig. S9). Finally, we note that, at ultra-high molecular blinking densities, the pixel-wise autocorrelation produces confidence levels for false localizations that rise above our chosen threshold (Fig. S8). A more effective approach in this situation may be to exploit the spatial correlation in the GradMap stack. Investigating an optimal strategy for leveraging correlations between multiple blinking events remains the subject of future studies.

## Methods

### Mathematical description of joint signal model

The proposed joint signal model together with a first-order approximation result in a linear forward model with a convex set of constraints on molecular parameters, denoted by 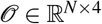^4^ (see Supplementary Information, section 2):

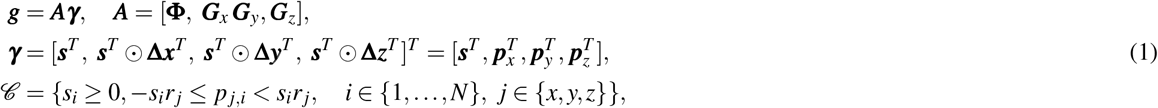

where *g* ϵ ℝ^*m*^ represents the vectorized noiseless image relayed by the microscope; 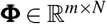 denotes the PSF matrix sampled at the grid points; 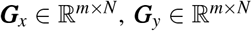, and 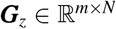 represent corresponding gradient matrices along *x*, *y*, and *z*, respectively; *N* is the number of object grid points; and *m* is the number of image pixels. Further, 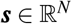 is the vectorized brightness and 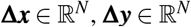, and 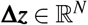 are the corresponding PSF gradient vectors (⊙ represents component-wise multiplication of two vectors) at each grid point. Additionally, 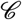 represents the convex set of constraints for a box centered at each grid point *i* with side lengths 2*r_x_*, 2*r_y_*, and *2r_z_* (Fig. 1(a)).

### Structured deconvolution

We develop a joint sparse deconvolution program to identify single molecules from their overlapping images. Our key insight is that the brightnesses *s* and position gradients 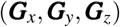 are jointly sparse. The structured deconvolution is cast as an optimization problem:

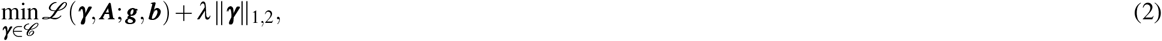

where 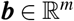 is the vectorized background and λ is a sparsity penalty parameter that balances the sparsity of the solution and likelihood of its observation. Further, 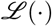(·) is the Poisson negative log likelihood given by

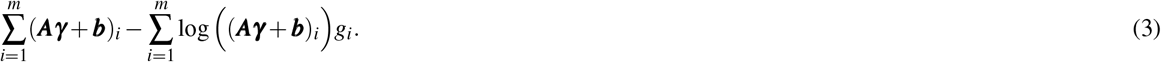

Finally, ‖ · ‖_1, 2_ denotes the mixed *l*_1,2_ norm, given by

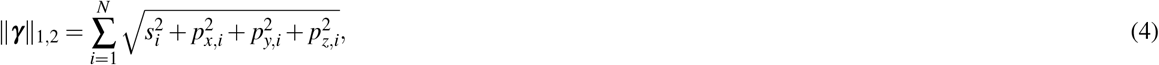

enforcing joint sparsity in 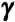^44^. The optimization problem given by (2) is a convex program (see Supplementary Information, section 3), and we develop a novel variant of the accelerated proximal gradient algorithm to efficiently solve it^45^. To calculate λ, we devised a computational tuning strategy (see Supplementary Information, section 3) accounting for each PSF 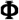.

The computational performance of FACON, SRRF, and RoSE-C are detailed in a supplementary table (Table S1).

## Data Availability

The source code and ground-truth molecule brightnesses and positions, simulated noisy images, and brightness and localization data output by RoSE and RoSE-C are available at: https://osf.io/vbptf/?view_only=5947calb9e8948bca2c70e536314d43d

## Acknowledgements

We thank Yoav Shechtman and W. E. Moerner for providing the tetrapod PSF and Tianben Ding, Oumeng Zhang, Eshan King, and Zhenqi Lu for their feedback on the manuscript. Research reported in this publication was supported by the National Science Foundation under grant number ECCS-1653777 and by the National Institute of General Medical Sciences of the National Institutes of Health under grant number R35GM124858 to M.D.L.

## Author contributions statement

H.M., A.N., and M.D.L. conceived the research; H.M. designed and wrote RoSE algorithm; H.M. analyzed simulated and experimental data with input from J.L. and M.D.L.; A.N. and M.D.L. supervised the research; all authors co-wrote and reviewed the manuscript.

## Additional information

**Competing financial interests**: The authors declare that they have no competing interests.

